# Genetic ablation of Sarm1 attenuates axonal degeneration in a mouse model of CMT2J

**DOI:** 10.64898/2026.10.28.762421

**Authors:** Anke Claessens, Roberta De Blasis, Cinzia Ferri, Serena Valenzano, Ubaldo Del Carro, Alessandra Mandelli, Roberto Furlan, Yo Sasaki, M. Laura Feltri, L. Wrabetz, Ghjuvan’Ghjacumu Shackleford, Maurizio D’Antonio

**Author notes:** Corresponding author: Maurizio D’Antonio. Neuroscience, School of Medical Sciences and Charles Perkins Centre, Faculty of Medicine and Health, University of Sydney, Sydney, Australia (Anke Claessens).

## Abstract

Charcot-Marie-Tooth (CMT) disease is the most common inherited neuropathy, affecting approximately 1 in 2,500 individuals, and comprises a clinically and genetically heterogeneous group of disorders involving more than 100 genes. Mutations in myelin protein zero (*MPZ*) typically cause the demyelinating form CMT1B, whereas the T124M substitution gives rise to CMT2J, a late-onset, predominantly axonal neuropathy with minimal myelin involvement. We previously generated a *Mpz*^T124M^ mouse model that recapitulates key features of the human disease, including progressive axonal degeneration and fibre loss in peripheral nerves. Because SARM1 (sterile α and TIR motif-containing protein 1) is a central executioner of programmed axonal degeneration, we crossed *Mpz*^T124M^ mice with *Sarm1*-null animals and assessed electrophysiological, morphological, and molecular parameters in both heterozygous and homozygous mutants.

*Sarm1* deletion did not rescue electrophysiological deficits in either genotype. However, in 12-month-old homozygous *Mpz*^T124M/T124M^ mice, *Sarm1* ablation partially reduced axonal degeneration in peripheral nerves and lowered plasma neurofilament light chain, a biomarker of neurodegeneration. In heterozygous *Mpz*^T124M/+^ mice, where axonal degeneration develops more slowly, *Sarm*1 deletion was associated with improved motor performance and reduced morphological signs of disease by 18 months of age.

Together, these findings indicate that SARM1 contributes to axonal degeneration in CMT2J and that its ablation confers partial protection, although it is insufficient to fully prevent disease progression, supporting SARM1 as a therapeutically relevant target in MPZ-associated axonal neuropathy despite incomplete disease rescue.

## INTRODUCTION

Axonal degeneration is a central pathological hallmark of peripheral neuropathies, and a common endpoint of diverse genetic, metabolic, and toxic insults. Metabolic disorders such as diabetes, and exposure to neurotoxic agents like chemotherapy, are among the leading causes of acquired axonopathies[1]. Inherited neuropathies, including Charcot-Marie-Tooth (CMT) disease, likewise converge on axonal degeneration, even when the primary mutation affects myelin proteins[2]. The loss of axonal integrity thus represents a shared mechanism driving functional decline across a broad spectrum of peripheral neuropathies.

Sterile alpha and TIR motif-containing protein 1 (SARM1) is a central executioner of active axonal degeneration and has emerged as a promising therapeutic target for neurodegenerative diseases[3]. Under normal conditions, SARM1 is present throughout the axon but kept inactive by the continuous activity of nicotinamide mononucleotide adenyltransferase 2 (NMNAT2), an axonal survival factor. NMNAT2 maintains low intracellular levels of nicotinamide mononucleotide (NMN) by catalysing its conversion to nicotinamide adenine dinucleotide (NAD^+^). This metabolic balance is critical, as NMN accumulation promotes SARM1 activation by binding to its allosteric pocket and inducing a conformational change, whereas NAD⁺ competes for the same site and stabilizes SARM1 in its inactive state. Upon axonal injury or NMNAT2 loss, the NMN:NAD⁺ ratio increases, activating the NADase activity of SARM1’s TIR domain[4–6]. Active SARM1 then cleaves NAD⁺ into nicotinamide (NAM) and adenosine diphosphate ribose (ADPR), leading to rapid depletion of NAD⁺ and ATP, metabolic collapse, and ultimately axonal degeneration[7,8].

Loss-of-function studies have demonstrated that SARM1 ablation confers robust protection across diverse models of axonal injury and disease. Axons from *dsarm*-deficient *Drosophila* or *Sarm1* knockout mice exhibit remarkable resistance to degeneration following axotomy[3]. Similarly, *Sarm1* deletion mitigates axonal loss in models of traumatic brain injury, chemotherapy-induced neuropathy, inherited neuropathies such as CMT2A and amyotrophic lateral sclerosis (ALS)[9–12]. SARM1 is an enzyme amenable to pharmacological modulation, and multiple strategies for SARM1 inhibition and targeted activation are rapidly developing[13]. These findings establish SARM1 as a master regulator of axonal degeneration and a highly promising therapeutic target for peripheral neuropathies and other disorders characterised by axonal loss.

Here, we investigated whether genetic ablation of *Sarm1* could attenuate disease progression in CMT2J mice. CMT is the most common inherited peripheral neuropathy, affecting 1 in 2500 people worldwide, and is characterised by progressive distal muscle weakness and sensory loss. CMT is broadly classified into demyelinating forms (CMT1) and axonal forms (CMT2), the latter being defined by primary axonal degeneration with limited demyelination.

CMT2J (also referred to as late-onset CMT1B) is caused by mutations in the myelin protein zero (*MPZ*) gene and represents a distinct phenotype among MPZ-associated neuropathies, as it displays predominantly axonal rather than demyelinating features. One of the best-characterised variants associated with CMT2J is the T124M mutation, resulting from a substitution of threonine to methionine at position 124. Notably, despite arising from a mutation in a myelin protein, CMT2J is marked by late-onset progressive axonal degeneration with only minimal signs of myelin abnormalities.

Our group previously generated and characterised an authentic model for CMT2J, carrying the *Mpz*^T124M^ mutation that recapitulates several features of the human disease[14]. Metabolic profiling of sciatic nerves from homozygous *Mpz*^T124M/T124M^ mice revealed a 55% reduction in NAD^+^ levels, accompanied by an increase in NAM and ADPR, consistent with activation of the SARM1 pathway. Given the central role of SARM1 in executing programmed axonal degeneration and its involvement in multiple models of peripheral neuropathy, we hypothesised that *Sarm1* deletion could preserve axonal integrity and modify disease progression in CMT2J. To test this hypothesis, we crossed *Mpz^T124M^* mice with *Sarm1* knockout animals and systematically evaluated functional, biochemical, and morphological markers of axonal degeneration.

## MATERIAL AND METHODS

### Animals

The *Mpz^T124M^* mouse model was previously generated and characterised by our group as described in Shackleford et al.[14]. In short, the mutation was targeted to a single *Mpz* allele by homologous recombination. For the experiments described in this paper, the line has been kept on the C57BL/6J background.

The *Sarm1^-/-^* line (B6.129X1-Sarm1tm1Aidi/J) was commercially acquired from The Jackson Laboratory (Strain #:018069)[15]. *Sarm1^-/-^* mice were crossed with *Mpz^T124M^* mice to generate the experimental *Mpz^T124M^Sarm1^-/-^* mice. For a more quantitative characterization of axonal damage, *Mpz^T124M^* mice were crossed with a *Thy1-YFP* reporter line (B6.Cg-Tg(Thy1-YFP)16Jrs/J). These transgenic mice were acquired from The Jackson Laboratory (Strain #:003709)[16] and are kept in hemizygosity.

Animals were housed in an environment with controlled temperature (21-24°C) and humidity, with a 12h light/dark cycle and *ad libitum* access to water and chow. All experiments performed on mice were conducted in accordance with experimental protocols approved by the San Raffaele Scientific Institute Animal Care and Use Committee (IACUC) and by the Italian Minister of Health.

### Genotyping

Genomic DNA was obtained from a piece of tail through phenol-chloroform extraction after incubation with a protein kinase lysis buffer. Day to day genotyping of *Mpz^T124M^* mutants was done with a PCR against the loxP sites. For *Thy1-YFP* and *Sarm1^-/-^* animals, the respective protocols from the Jackson Laboratory were used.

### Electrophysiology and behavioural analysis

Electrophysiology was performed as previously described by Della-Flora Nunes et al.[17]. At the ages indicated in the text, mice were put under light anaesthesia with 0.02 mL/g of bodyweight tribromoethanol (Avertin; Sigma-Aldrich) and placed under a heating lamp. With an electromyographer (Neuro-MEP-Micro; Neurosoft), peripheral nerves were stimulated with short electrical pulses, and the subsequent muscle responses were recorded. The sciatic nerve conduction velocity (NCV) was obtained with steel monopolar needle electrodes. One pair of stimulating electrodes was inserted subcutaneously near the nerve at the ankle, a second pair of electrodes was placed at the sciatic notch, and a third pair over the dorsum of the spine. Compound muscle action potential was recorded with an active electrode inserted in muscles in the middle of the paw and a reference needle in the skin between the first and second digits. Sciatic nerve F wave latency measurement was obtained by stimulating the nerve at the ankle and recording the responses in the paw muscle, with the same electrodes employed for the NCV study. For the electrophysiology studies, a minimum of *n* = 5 mice per genotype were used and resulted in two measurements (= sciatic nerve of both the left and right hind paw) per mouse, unless stated otherwise. Asymmetrical measurements were excluded.

Muscle endurance was tested through the inverted grid test. Mice were positioned on a wired grid, placed on an elevated support. The grid was then inverted and the time that the mice were able to hold on was measured. Mice were assessed in random order, on two separate days with three trials per day. All six trials were averaged.

### Morphological analysis

Transgenic mice were sacrificed at the indicated ages and sciatic and femoral nerves were dissected and fixed in 2% glutaraldehyde in phosphate buffer, osmicated in 1% OsO4, alcohol dehydrated, infiltrated with propylene oxide, and embedded in Epon. Transverse semithin sections (1 µm thick) were stained with toluidine blue and acquired with a Leica DM5000 microscope equipped with a DFC480 digital camera. Pictures were taken with a 100x objective, and PTGui software v.10 (New House Internet Services BV) was used to reconstruct a complete image of the nerve. These reconstructions were then used to measure the circumference of the nerve, the total number of myelinated fibres and degenerative axons. We used ImageJ Fiji v1.52p software and analysed a minimum of 3 reconstructed nerve images per genotype. Ultrathin sections were stained with lead citrate and photographed with a Zeiss (Oberkochen, Germany) EM10 electron microscope.

### Simoa plasma NfL measurement

Blood was collected from the submandibular (facial) vein of alive but anesthetised mice (*n* = 3 per genotype) and transferred in EDTA coated tubes. Samples were incubated on ice for 15 minutes and centrifuged for 20 min at 3000 rpm at 4°C. The supernatants (plasma) were transferred into new tubes and stored at -80°C until analysis. Plasma neurofilament light chain concentrations were measured using the Simoa® NfL assay as described by Sandelius et al. (2018)[18].

### Metabolite measurement

Metabolite extraction from sciatic nerves was performed and measured as described by Sasaki et al. (2020)[19]. In brief, nerves were snap-frozen in liquid nitrogen, homogenised by sonication in 50% MeOH/water, and cleared by centrifugation (13,000 *g*, 10 min, 4°C). Following chloroform-based phase separation, the aqueous fraction was lyophilised and then reconstituted in 5 mM ammonium formate. NAD⁺ and ATP were quantified by LC-MS/MS using a C18 reverse-phase column (Atlantis T3; Waters) equipped with HPLC system (Agilent 1290 LC) and coupled to a triple quadrupole mass spectrometer (Agilent 6470 MassHunter; Agilent) under positive ESI-MRM mode. Metabolites were normalised to protein content determined by BCA assay (Pierce^TM^; Thermo Scientific).

### Teased nerve preparation

Sciatic nerves from *Thy1-YFP* mice were freshly dissected, fixed in 4% paraformaldehyde for 30 min, washed in 1x PBS and then teased immediately. The epineurium was removed with dissecting forceps so that the entire nerve could be whole mounted with VECTASHIELD® (H-1000-10, VectorLabs) onto glass adhesion microscope slides (Epredia). Slides were allowed to dry for at least 1h and stored at -80 °C until use.

A minimum of 5 images per nerve were taken randomly but along the entire teased sciatic nerve with a confocal Leica SP5II microscope and analysed using ImageJ Fiji v1.52p. The level of axonal degeneration was measured by counting the number of swellings and fragmentations over the total number of THY1-YFP positive fibres in that field and then averaged per nerve.

### Statistical analyses

Data collection and analysis were performed blinded to the genotype. All statistical analyses were conducted using GraphPad Prism 9.01 and significance was assessed by Student’s t test or one-way ANOVA followed by a Tukey’s post hoc test correction. A *p*-value ≤ 0.05 was considered statistically significant.

## RESULTS

### *Sarm1* ablation provides partial protection from CMT2J disease progression

To investigate whether SARM1 contributes to CMT2J pathogenesis, we crossed *Mpz*^T124M^ mice with *Sarm1* knockout mice (*Sarm1^-/-^*)^15^. Male and female offspring of all genotypes (wild type; WT, *Mpz*^T124M/+^ or *Mpz^T124M/T124M^* with or without *Sarm1* deletion) were subsequently evaluated for functional and pathological outcomes.

Our group previously showed that *Mpz*^T124M^ mice display progressive neurophysiological defects starting at 6 months of age. To assess early presymptomatic effects of SARM1 ablation, like delayed disease onset or slowed disease progression, we analysed the first cohort at 4 months old. While no significant alterations in nerve conduction velocity (NCV) could be detected, we observed mild electrophysiological changes such as a trend towards reduction in compound muscle action potentials (CMAPs), and an increase of F wave latency in homozygous *Mpz*^T124M/T124M^ mice. *Sarm1* deletion did not ameliorate these defects at this early timepoint (**Figure S1A-C)**.

Peripheral nerves from *Mpz^T124M^Sarm1^-/-^* and control mice were then dissected for morphological and morphometric analyses. Consistent with previous reports indicating that SARM1 is dispensable for myelination, *Sarm1* deletion did not affect WT sciatic nerve morphology[20]. Furthermore, *Mpz^T124M^* sciatic nerves looked completely normal as well (**Figure S1D**).

Because patients with CMT2J develop prominent motor disability and weakness[21], we also examined femoral nerves that, differently from sciatic nerves, allow separate analysis of motor and sensory branches. In contrast to the sciatic nerves, quadriceps motor nerves (QN) from homozygous *Mpz^T124M/T124M^* mice already displayed a severe phenotype at 4 months, including extensive axonal loss but also demyelination and onion bulbs formation (**Figure 1A**). Notably, the predominantly sensory saphenous branch was unaffected (**Figure S1E**). Although *Sarm1* deletion did not fully rescue the severe degenerative morphology of QN nerves, the quantification of nerve fibre density revealed a partial protective effect. In fact, whereas fibre density was significantly reduced in *Mpz^T124M/T124M^* nerves, values in *Mpz^T124M/T124M^Sarm1^-/-^* mice were comparable to WT (**Figure 1B-C**). These findings indicate that *Sarm1* deletion partially ameliorates early structural pathology in motor nerves, despite the absence of electrophysiological improvement.

**Figure 1:**
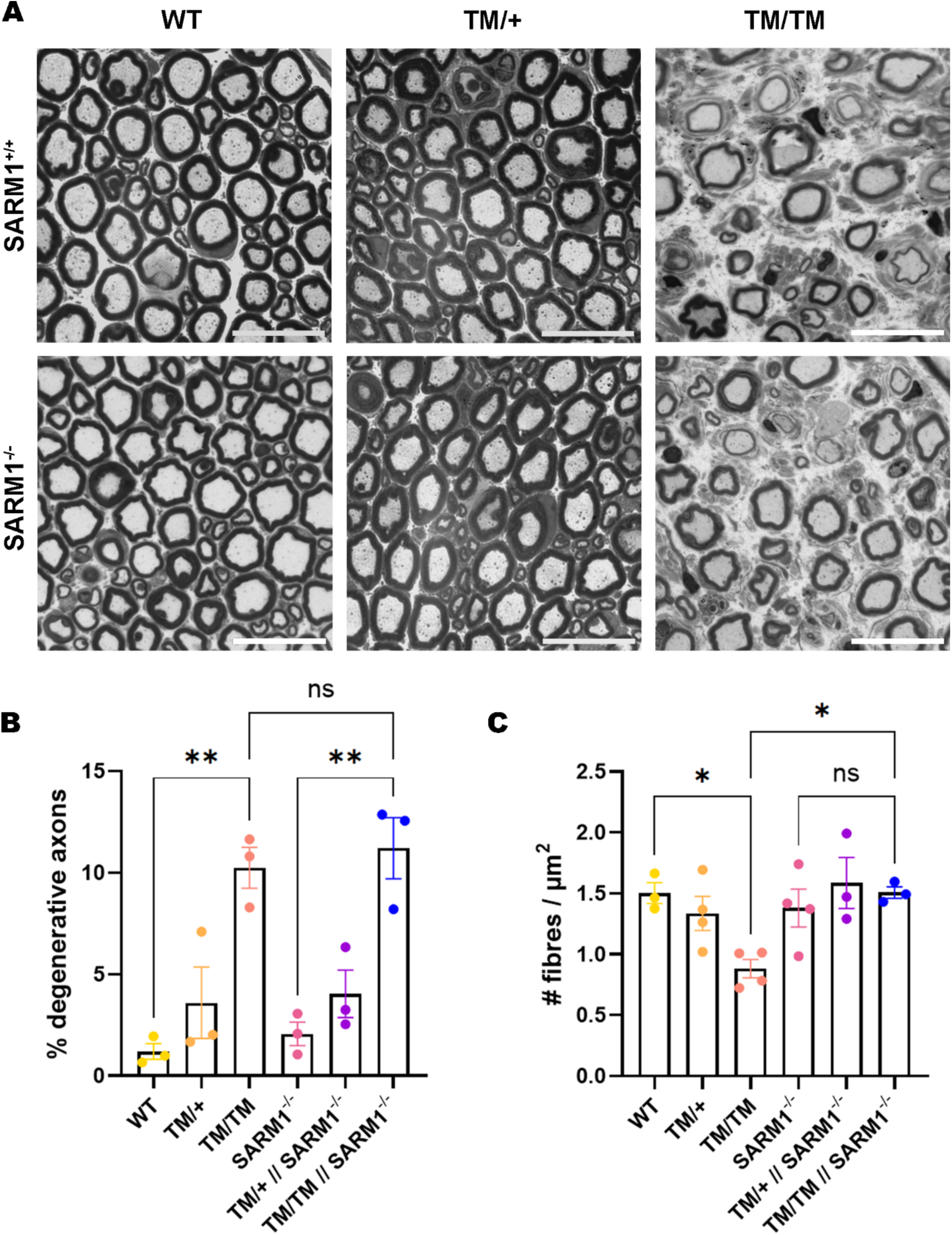
Femoral nerve morphology of 4-month-old Mpz^T124M^Sarm1^-/-^ mice. (A) Representative semithin sections of femoral nerves from 4-month-old mice. (B) Quantification of degenerative axons in 4-month-old femoral nerves and (C) quantification of the density of myelinated fibres in femoral nerves. Scale bars = 20µm; n = 3-4 nerves per genotype. *p<0.05 by one-way ANOVA with Tukey’s post hoc. Graphs indicate means ± SEM.

By 8-months of age, *Mpz^T124M^* mice exhibited a phenotype consistent with disease progression. The electrophysiological trend differences observed at 4 months had now reached statistical significance, with homozygous *Mpz^T124M/T124M^* mice displaying reduced CMAP amplitudes and increased F wave latencies. However, these abnormalities were not ameliorated by *Sarm1* deletion (**Figure 2A-C**). Persistent degeneration was observed in QN of both *Mpz^T124M^* and *Mpz^T124M^Sarm1^-/-^* mice, comparable to that observed at 4 months (**Figure S2)**. Notably, pathological alterations started to become evident also in sciatic nerves of homozygous *Mpz^T124M/T124M^* mice (**Figure 2D**). Quantifications showed a trend towards increased axonal degeneration and reduced fibre density, neither of which was measurably affected by Sarm1 ablation, indicating that SARM1 loss does not modify disease progression at this stage (**Figure 2E-F**).

**Figure 2:**
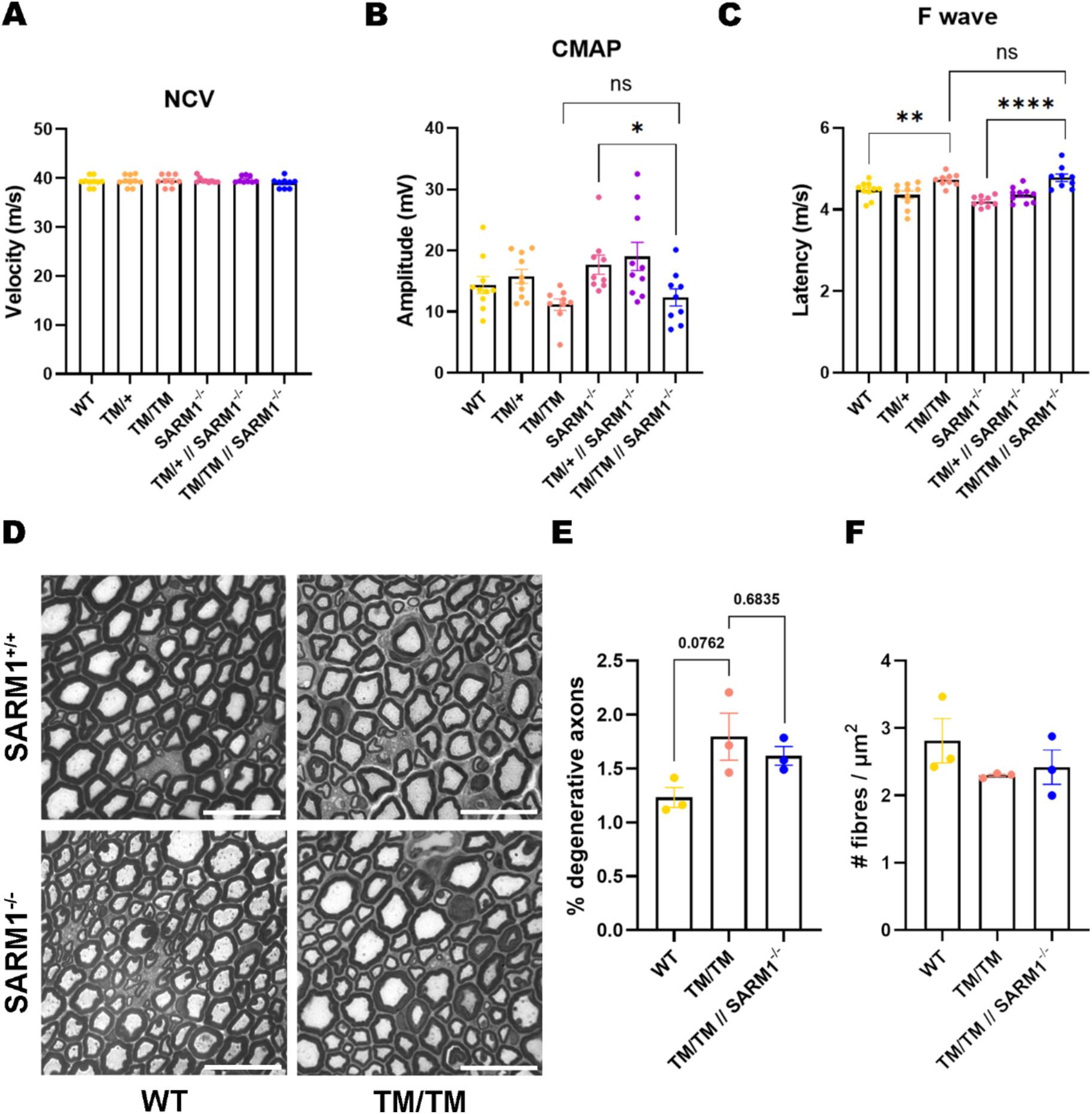
Electrophysiological parameters and sciatic nerve morphology show mild neuropathy in 8-month-old Mpz^T124M^ and Mpz^T124M^Sarm1^-/-^ mice. (A) Nerve conduction velocity (NCV), (B) amplitude of compound muscle action potentials (CMAPs) and (C) latency of the F wave were measured at 8 months of age. n = 5 mice per genotype. (D) Semithin sections of sciatic nerves obtained from 8-month-old mice with the quantification of degenerative axons (E) and fibre density (F) in homozygous Mpz^T124M/T124M^ and Mpz^T124M/T124M^ Sarm1^-/-^ mice. Scale bars = 20µm. ****p<0.0001; **p<0.01; *p<0.05 by one-way ANOVA with Tukey’s post hoc. Graphs indicate means ± SEM.

Consistent with previous reports[14], electrophysiological analysis at 12 months showed significantly reduced NCV and CMAP amplitudes, together with increased F wave latency in homozygous *Mpz*^T124M/T124M^ mice (**Figure 3A-C**). None of these parameters however were changed by *Sarm1* deletion. As previously described[14], sciatic nerves from *Mpz^T124M/T124M^* mice showed an increased number of degenerative fibres at this time point, despite largely preserved myelin compaction. Although slightly reduced in numbers, degenerative axons were also observed in *Mpz^T124M/T124M^Sarm1^-/-^* mice, (**Figure 3D-E**). In contrast, analysis on QN nerves revealed a more severe pathological phenotype in *Mpz^T124M/T124M^* mice which was partially ameliorated by SARM1 deletion (**Figure 3F-G**). This suggest that *Sarm1* deletion confers nerve-specific protection of axonal degeneration, despite failing to improve electrophysiological outcomes.

**Figure 3:**
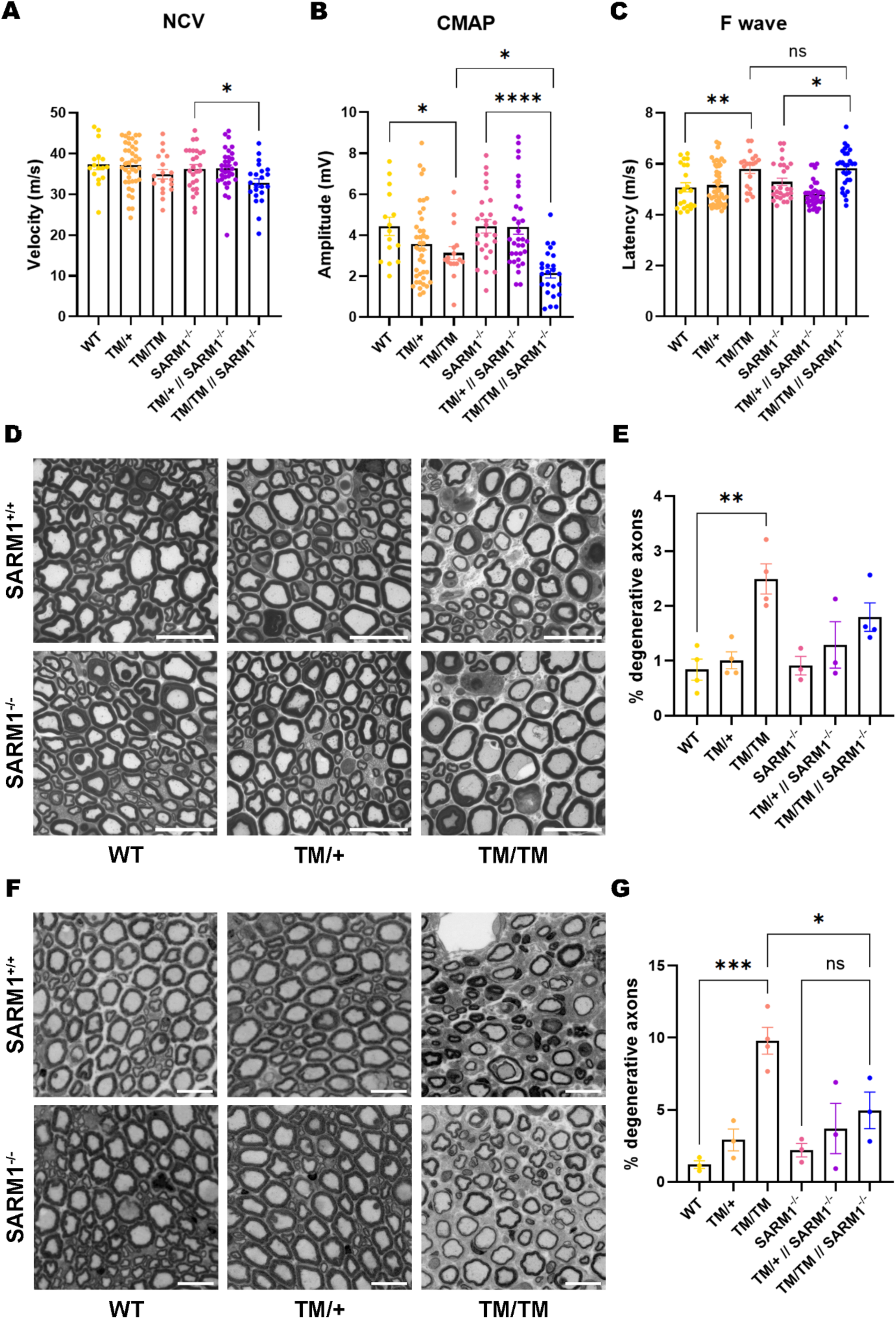
Sarm1 deletion differentially affects axonal degeneration in sciatic and femoral nerves of 12-month-old Mpz^T124M^ mice. (A) Nerve conduction velocity (NCV), (B) compound muscle action potentials (CMAPs) amplitude and (C) F-wave latency were measured at 12 months of age. n > 10 mice per genotype. (D) Representative semithin sections from sciatic nerves and (E) quantification of degenerating fibres. (F) Representative semithin sections of 12-month-old femoral motor nerves and (G) quantification of degenerating fibres. Scale bars = 20µm; n = 3-4 nerves per genotype for D-G. ****p<0.0001; ***p<0.001; **p<0.01; *p<0.05 by one-way ANOVA with Tukey’s post hoc. Graphs indicate means ± SEM.

### Sensitive detection of axonal degeneration reveals a protective effect of *Sarm1* deletion

To obtain a more sensitive and quantitative assessment of axonal degeneration, we crossed *Mpz^T124M^* and *Mpz^T124M^Sarm1^-/-^* mice with a transgenic line expressing yellow fluorescent protein (YFP) under the control of regulatory elements of the mouse *Thy1* gene (*Thy1-YFP*)[16]. THY1 is a glycoprotein expressed on the surface of neuronal and glial cells[22], enabling robust fluorescent labelling of both motor and sensory neurons and allowing direct visualisation and quantification at single-fibre resolution. Teased sciatic nerves from 12-month-old triple transgenic mice were imaged by confocal microscopy (**Figure 4A**). Axonal degeneration was quantified by scoring characteristic degenerative features such as focal axonal swellings and fragmented fibres resembling those seen during Wallerian degeneration and normalizing these degeneration events over the total number of THY1-YFP positive fibres. This quantification showed that degenerative axons were increased in *Mpz^T124M^* animals in comparison to WT, and that *Sarm1* deletion markedly reduced axonal degeneration in both heterozygous and homozygous mutants. In homozygous *Mpz^T124M/T124M^* mice, the proportion of degenerating axons was reduced by approximately 50% following *Sarm1* ablation, while *Mpz^T124M/+^Sarm1^-/-^* mutants displayed levels of axonal degeneration comparable to WT and *Sarm1^-/-^* mice (**Figure 4B**). This higher-resolution assessment of axonal integrity demonstrates that *Sarm1* significantly attenuates axonal degeneration in CMT2J mice, consistent with a direct contribution of SARM1 to axonal pathology in this disease.

**Figure 4:**
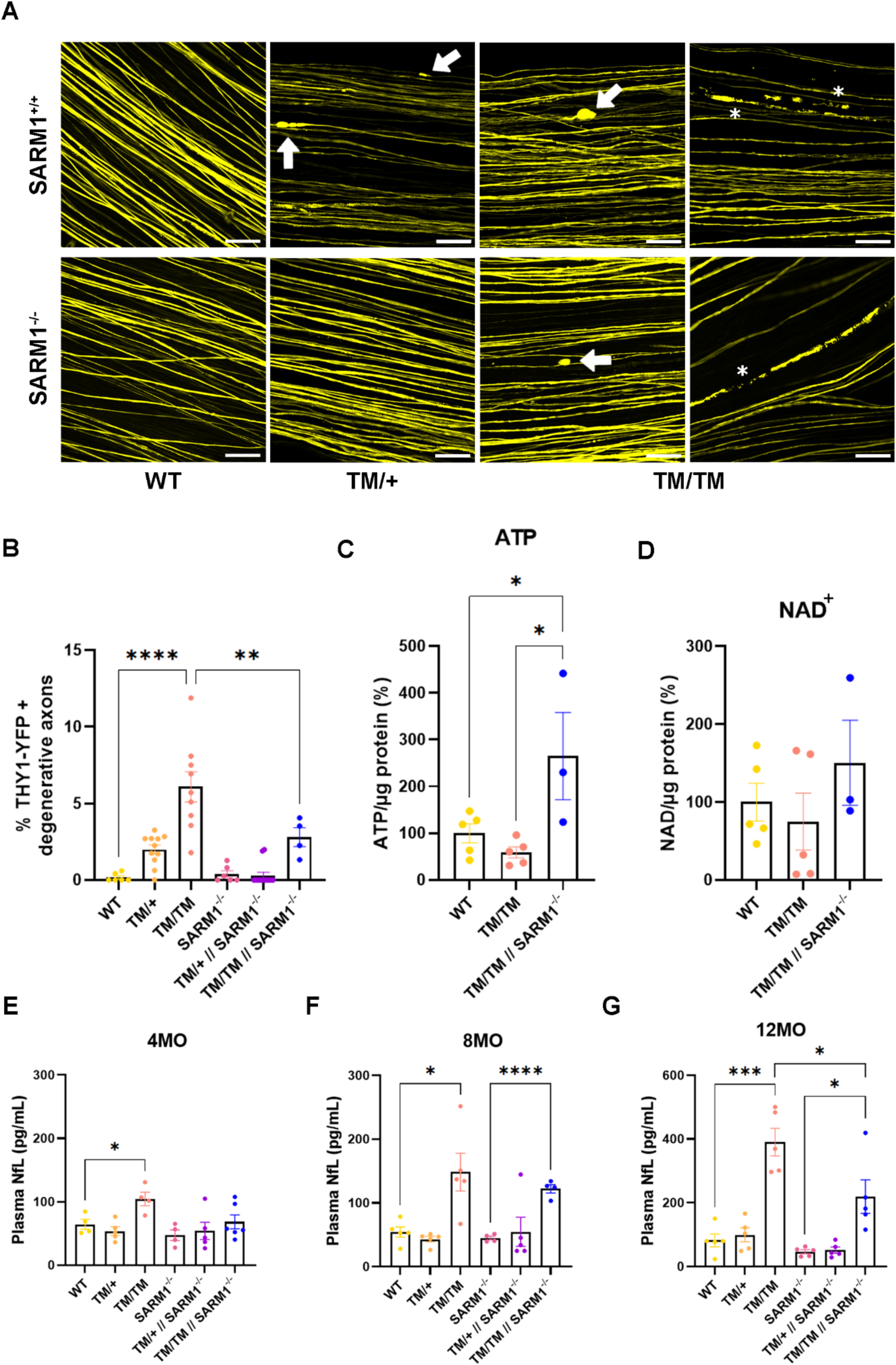
Sarm1 deletion attenuates axonal degeneration and metabolic and biochemical markers of disease in Mpz^T124M^ mice. (A) Representative confocal images of teased fibres from 12-month-old Mpz^T124M^ and Mpz^T124M^ Sarm1^-/-^ mice in a Thy1-YFP background. Arrows indicate degenerating axons; asterisks indicate axonal fragmentation. Scale bars = 100 µm. (B) Quantification of degenerating axons in teased sciatic nerve fibres. n = 4-13 nerves per genotype. ****p<0.0001; **p<0.01; *p<0.05 by one-way ANOVA with Tukey’s post hoc test. (C) ATP and (D) NAD^+^ levels were measured in sciatic nerves from 12-month-old mice via LC-MS/MS. n = 3-5 nerves per genotype. *p<0.05 by one-way ANOVA with Tukey’s post hoc test. (E-G) Plasma neurofilament light chain (NfL) measurements from (E) 4, (F) 8 and (G) 12-month-old mice. n = 4-5 mice per genotype. ****p<0.0001; **p<0.01 by Student’s t test. All graphs indicate means ± SEM.

### *Sarm1* ablation partially normalizes nerve metabolites and degeneration biomarker levels

To provide further evidence that the protective effects observed in *Mpz^T124M^* mice were associated with suppression of SARM1-dependent pathways, we assessed metabolic alterations previously identified in sciatic nerves from 12-month-old *Mpz^T124M/T124M^* mice. Using mass spectrometry, we measured ATP and NAD^+^ levels in WT, *Mpz^T124M/T124M^* and *Mpz^T124M/T124M^Sarm1^-/-^* nerves. As reported previously, *Mpz^T124M/T124M^* nerves exhibited reduced ATP and NAD^+^ levels, consistent with the metabolic signature of SARM1 activation[14]. Despite considerable internal variability, both metabolites displayed a trend toward normalization following *Sarm1* deletion (**Figure 4C-D**) supporting the notion that SARM1 contributes to metabolic abnormalities associated with CMT2J.

Next, we examined plasma neurofilament light chain (NfL), a well-established biomarker for neurodegeneration[18] that is progressively elevated in *Mpz^T124M^* mice[14]. Blood was sampled from *Mpz^T124M^* and *Mpz^T124M^Sarm1^-/-^* mice at 4, 8 and 12 months of age. A small increase of NfL concentration was already detectable in plasma samples from 4-month-old *Mpz^T124M/T124M^* mice, which became evident and significant at 8 months (**Figure 4E-F**). Although NfL concentrations also increased with age in plasma from *Mpz^T124M/T124M^Sarm1^-/-^* mice, the magnitude of the increase was consistently reduced. Interestingly, by 12 months of age there was a 6-fold increase of NfL levels in *Mpz^T124M/T124M^* compared to WT mice while SARM1 deficient *Mpz^T124M/T124M^* mice only showed a two-to three-fold increase (**Figure 4G**). Together, these findings confirm that *Sarm1* deletion attenuates both the metabolic and biochemical consequences of axonal damage in CMT2J mice. Combined with the observed reduction in THY1-YFP degenerating axons, these results provide strong evidence that SARM1 contributes to axonal degeneration in CMT2J mice.

### SARM1 deletion preserves motor function and attenuates axonal pathology in aged _Mpz_T124M/+ _mice_

Notably, the protective effects of *Sarm1* deletion observed in homozygous *Mpz^T124M/T124M^* mice were most evident using sensitive measures of axonal degeneration, while standard electrophysiological and morphological analyses revealed only modest improvements. Because heterozygous *Mpz^T124M/+^* mice develop an even more slowly progressive neuropathy that closely resembles the human disease, we investigated whether the impact of *Sarm1* deletion would become more apparent at advanced stages of disease progression. Therefore, electrophysiology and grip strength were assessed in a small cohort of 18-month-old *Mpz^T124M/+^* and *Mpz^T124M/+^Sarm1^-/-^* mice (**Figure 5A-C**). Electrophysiological parameters were slightly affected in all animals, including WT mice. This decline is most likely due to their advanced age, as studies have shown that both NCV and muscle action potential amplitude progressively decrease with age[23,24]. No clear genotype-dependent differences were observed. In contrast, the inverted grid test revealed a marked reduction in *Mpz^T124M/+^* mice, that could not hold on for more than 10 seconds. Despite the small cohort size, *Mpz^T124M/+^Sarm1^-/-^* mice consistently outperformed *Mpz^T124M/+^* littermates, on average reaching values comparable to WT controls (**Figure 5D**).

**Figure 5:**
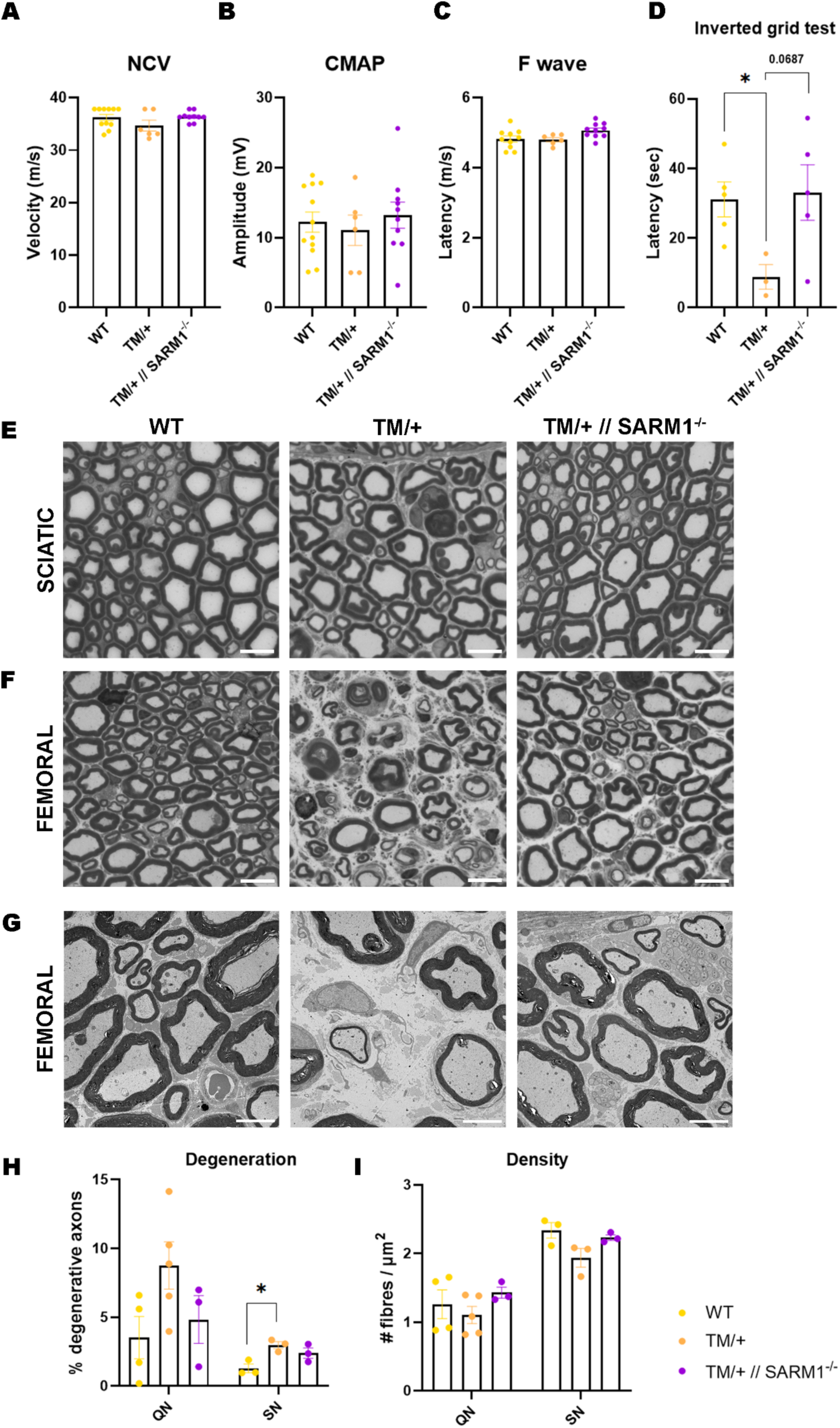
Sarm1 improves motor performance and attenuates axonal pathology in aged Mpz^T124/+^ mice. (A) Nerve conduction velocity (NCV), (B) compound muscle action potentials (CMAPs) amplitudes (C) F-wave latency and (D) inverted grid test were measured at 18 months of age. n = 3-5 mice per genotype. *p<0.05 by one-way ANOVA with Tukey’s post hoc. Representative semithin sections of (E) sciatic nerves (SN) and (F) quadriceps femoral nerves (QN) of Mpz^T124M/+^ mice. (G) Representative electron micrographs of QN. Scale bars = 20 µm. (H) Quantification of degenerative fibres in QN and SN nerve fibres. (I) Quantification of myelinated fibre density in QN and SN fibres. *p<0.05 by Student’s t test. Graphs indicate means ± SEM.

Recent work from Klein et al. showed that heterozygous *Mpz^T124M/+^* mice develop a severe axonopathy at 18 months of age[25]. Consistent with these observations, semithin sections revealed prominent pathological changes in both sciatic and femoral nerves, including degenerating fibres, regenerative clusters, axonal loss, and secondary demyelination, with the femoral nerve displaying the most severe phenotype **(Figure 5E-F)**. Ultrastructural EM analysis further confirmed extensive axonal degeneration, characterised by abundant cellular debris, regenerative clusters, and structures resembling denervated Schwann cells or dysfunctional Büngner cells[26] (**Figure 5G** and **Figure S3)**. Notably, these pathological changes appeared less pronounced in *Mpz^T124M^*^/+^Sarm1^-/-^ nerves. Quantitative analysis of semithin sections revealed increased fibre density in both sciatic and femoral nerves and a reduction of degenerative fibres in femoral nerves of *Sarm1*-deficient animals (**Figure 5H-I**). Although based on a limited number of aged animals, the concordant behavioural and morphological improvements point consistently to a protective role of *Sarm1* ablation in heterozygous *Mpz^T124M/+^* mice. Together with the reduced axonal degeneration, plasma NfL and normalised metabolic markers in in homozygous *Mpz^T124M/T124M^* mice, these findings reinforce a contribution of SARM1 to axonal degeneration in CMT2J.

## DISCUSSION

Axonal degeneration is a central feature of CMT2J, and evidence from our previous study suggests that SARM1 activation may contribute to the underlying pathomechanism[14]. Given the observed reduction of NAD⁺ in homozygous *Mpz*^T124M/T124M^ mice, which is a hallmark of SARM1-dependent degeneration, we investigated whether genetic deletion of *Sarm1* could mitigate disease progression. To this end, *Mpz*^T124M^ mice were crossed with *Sarm1* null animals, and the impact of SARM1 loss on electrophysiological, morphological, and molecular hallmarks of neuropathy was systematically evaluated.

Genetic ablation of *Sarm1* did not prevent or delay the electrophysiological impairments that characterise CMT2J. Motor conduction velocity and compound muscle action potentials remained reduced and F wave latency was still prolonged in *Mpz*^T124M^*Sarm1^⁻/⁻^* mice at all tested ages. Similarly, semithin sections of sciatic nerves showed persistent signs of degeneration without obvious structural rescue. However, morphological quantifications indicate higher preservation of total axon numbers in nerves of *Mpz*^T124M^*Sarm1^⁻/⁻^* mice, pointing towards delayed axon death. Additionally, more sensitive analyses of teased sciatic nerves using the *Thy1-YFP* reporter line revealed a significant reduction in axonal fragmentation in *Mpz*^T124M^*Sarm1^⁻/⁻^* mice, indicating partial morphological protection. This underscores the value of high-resolution approaches for detecting dynamic axonal degeneration that may not be evident in static histological sections.

Interestingly, the extent of protection varied across peripheral nerves. While pathological changes in the sciatic nerve of homozygous *Mpz*^T124M/T124M^ mice become evident only from 8 months of age, the femoral quadricep nerve, predominantly composed of motor fibres, exhibited early and severe pathology including axonal loss, demyelination, and onion bulb formation from as early as 4 months. In this nerve, *Sarm1* deletion resulted in increased myelinated fibre density and reduced degeneration, indicating a degree of selective protection and delayed degeneration. In contrast, the predominantly sensory saphenous nerve appeared unaffected in *Mpz*^T124M^ mice. This pattern may reflect a difference in vulnerability between motor and sensory axons, potentially influenced by factors such as axon length or trophic support. Such selectivity is consistent with the clinical presentation of CMT2J, in which motor symptoms are predominant despite the presence of sensory abnormalities [27,28]. Nerve-specific vulnerability is a common feature across peripheral neuropathies, although its direction is highly disease-dependent. This is exemplified in CMT, where different mutations within the same gene can cause either a sensorimotor or a purely motor neuropathy[29]. The specific mechanism underlying this nerve-specific vulnerability remains unresolved[30].

Consistent with the more sensitive morphological findings, plasma NfL levels were significantly elevated in aging *Mpz*^T124M/T124M^ mice but were consistently lower in the SARM1 deficient animals. At 12 months of age, *Sarm1* deletion reduced the NfL increase from six-fold to approximately two-to three-fold relative to WT, supporting a partial attenuation of axonal degeneration. This is consistent with findings from a model of chemotherapy-induced peripheral neuropathy, in which SARM1 inhibition reduced circulating NfL levels and protected axon structure and function[31]. These observations suggest that the axonal degeneration and the associated release of neurofilament into the circulation in *Mpz^T124M^* mice are at least partially mediated by SARM1-dependent mechanisms.

In our study, we analysed both hetero-and homozygous *Mpz^T124M^* mice to assess potential differences in the protective effect of *Sarm1* deletion. Importantly, most CMT2J patients carry a single copy of the mutant *Mpz^T124M^* allele, although rare homozygote cases have been reported. They typically present with earlier onset and more severe symptoms than heterozygous patients[32], consistent with our observations in the corresponding mouse models. A recent study from Klein et al. described extensive axonopathy in 18-month-old heterozygous *Mpz^T124M/+^* mice, particularly in femoral nerves, including typical pathological changes such as periaxonal vacuoles and regenerative clusters[25]. We observed comparable alterations in our cohort, and ultrastructural analysis revealed denervated Schwann cells with fragmented myelin and cellular protrusions. These features are consistent with a primary axonal neuropathy in which Schwann cells persist after axonal loss but remain structurally disorganised. In contrast, nerves from *Mpz^T124M/+^Sarm1^-/-^* mice were markedly less affected, with reduced axonal loss and less pronounced myelin defects in both sciatic and quadriceps nerves. This was accompanied by improved muscle strength in 18-month-old *Mpz^T124M/+^Sarm1^-/-^* mice, suggesting that *Sarm1* deletion may help preserve axonal integrity and function at advanced disease stages in this model.

Notably, this partial rescue closely parallels the results of the Klein et al. study, where CSF1R-mediated macrophage depletion significantly ameliorated disease features[25]. Consistent with this, re-examination of the femoral nerve semithin sections from our 18-month-old *Mpz^T124M/+^Sarm1^-/-^* cohort revealed fewer phagocytosing macrophages (**Figure S4**). This observation is qualitative and will require systematic quantification, ideally by electron microscopy, but it points towards a link between SARM1 activity and neuroinflammation.

This idea is consistent with findings in a congenital NMNAT2-deficiency model, in which chronic SARM1 activation drove macrophage infiltration and macrophage depletion substantially ameliorated the neuropathy[33]. Together, these observations are consistent with a conserved SARM1–macrophage degenerative axis in chronic neuropathies, in which SARM1-dependent axonal degeneration promotes macrophage recruitment and activation that may in turn amplify disease progression. In this scenario, targeting SARM1, macrophages, or both could offer therapeutic benefit where chronic axonal degeneration and neuroinflammation co-exist. Further delineation of this pathway will be important to resolve the interplay between neuronal-intrinsic and immune-mediated mechanisms of disease progression.

Although SARM1 loss did not fully rescue the CMT2J phenotype, the improvements observed in axonal morphology, circulating NfL levels, and motor performance are noteworthy. Similar partial or context-dependent effects of SARM1 ablation have been reported in other neuropathy models. For example, in a mouse model of sporadic ALS, *Sarm1* deletion modified biomarkers of neurodegeneration without altering disease progression[34]. Similarly, *Sarm1* deletion failed to improve the phenotype of a CMT1A mouse model[35]. However, this model is primarily characterised by dysmyelination with little overt axonal degeneration, making a robust benefit from SARM1 ablation unlikely from the outset. More recently, *Sarm1* deletion or inhibition was evaluated across six different mouse models of CMT but failed to improve neuropathic phenotypes despite a reduction in plasma NfL[36,37]. Importantly, these mouse lines were selected based on their clinical relevance rather than on evidence of SARM1 pathway activation. Together with our findings, these studies suggest that the therapeutic efficacy of SARM1 inhibition may depend on the extent to which SARM1 contributes to the underlying disease mechanism.

Consistent with this concept, a recent study in CMT2A rats showed robust protection following *Sarm1* deletion with rescue of multiple hallmarks of the disease[11]. These results collectively underscore that SARM1 activation is not a universal driver of pathology but rather contributes to disease progression in a context-dependent manner. The greatest benefits of SARM1 loss have been observed in models characterised by clear metabolic or mitochondrial dysfunction, as in mitofusin 2 (MFN2)-associated CMT2A, supporting the concept that neuropathies with evidence of SARM1 pathway activation, such as NAD⁺/NMN imbalance, are more likely to benefit from SARM1-targeted therapies. Accordingly, homozygous *Mpz^T124M/T124M^* mice displayed a marked reduction in ATP and NAD⁺ and an accumulation of NAM and ADPR[14], providing biochemical evidence of SARM1 activation. Importantly, *Sarm1* deletion restored ATP and NAD^+^ levels in CMT2J peripheral nerves, demonstrating effective suppression of the underlying metabolic defect.

## CONCLUSION

Taken together, our findings establish that SARM1 contributes to axonal degeneration in CMT2J. Genetic deletion of *Sarm1* normalised nerve ATP and NAD⁺, reduced axonal fragmentation and circulating NfL, and preserved motor function at advanced disease stages, confirming that part of the axonal pathology in this model is mediated through SARM1-dependent Wallerian-like degeneration. At the same time, SARM1 loss did not rescue the electrophysiological deficits or halt overall disease progression, indicating that SARM1 activation is likely one of several converging mechanisms driving the neuropathy rather than its sole cause. The protection was also nerve-type-dependent, being most evident in severely affected motor nerves, suggesting that the benefit of targeting SARM1 is greatest in nerves with the highest burden of axonal degeneration.

These results carry two implications for the field. First, they add CMT2J to the growing but still selective list of neuropathies in which SARM1 is demonstrably involved in the pathology, and they reinforce an emerging principle: the therapeutic value of SARM1 inhibition is not universal but depends on evidence of pathway activation, such as the NAD⁺/NMN imbalance we observe here. This supports a mechanism-guided approach to therapeutic stratification, in which metabolic or biomarker signatures of SARM1 engagement could help identify the axonal neuropathies most likely to respond. Second, because SARM1 deletion conferred measurable but incomplete neuroprotection, our data indicate that SARM1 inhibition is unlikely to be sufficient as a standalone therapy. Instead, it should be considered as a key component of future combination-based therapeutic strategies targeting complementary pathogenic mechanisms. Such an approach may be particularly relevant for CMT2J and other axonal neuropathies for which no disease-modifying treatment currently exists.

## LIST OF ABBREVIATIONS

ADPR: Adenosine diphosphate ribose
ALS: Amyotrophic lateral sclerosis
ATP: Adenosine triphosphate
CMAP: Compound muscle action potential
CMT: Charcot-Marie-Tooth
MFN2: Mitofusin 2
MPZ: Myelin protein zero
NAD^+^: Nicotinamide adenine dinucleotide
NAM: Nicotinamide
NCV: Nerve conduction velocity
NFL: Neurofilament light chain
NMN: Nicotinamide mononucleotide
NMNAT2: Nicotinamide mononucleotide adenyltransferase 2
QN: Quadriceps nerve
SARM1: Sterile α and TIR (Toll-like interleukin 1 receptor) motif-containing protein 1
SN: Sciatic nerve
TM: T124M Threonine by methionine substitution at position 124 of MPZ
WT: Wild type
YFP: Yellow fluorescent protein

## DECLARATIONS

## Ethics approval and consent to participate

All experiments performed on mice were conducted in accordance with experimental protocols approved by the San Raffaele Scientific Institute Animal Care and Use Committee (IACUC) and by the Italian Minister of Health.

## Consent for publication

Not applicable

## Availability of data and material

All the data that was produced by this study is available in the manuscript and the supplementary information.

## Authors’ contributions

GGS and MD designed the study. MD, GGS, MLF and LW raised funds for the study. AC carried out the experiments and performed the formal data analysis. RDB contributed to mouse colony maintenance, motor studies and peripheral nerve collection. CF prepared nerve sections for imaging. SV and UDC performed electrophysiology. GGS performed the THY1-YFP imaging and quantification of the 12MO cohort. AM and RF carried out the NfL measurements. YS ran the metabolomic analysis. AC drafted the original manuscript, MD reviewed and edited the final version of the manuscript, and all authors read and approved the final version of the manuscript.

## Acknowledgements

We thank Prof. Aaron DiAntonio for critical reading of the manuscript. We thank the Alembic facility at the San Raffaele Scientific Institute for excellent imaging help. We would also like to thank and acknowledge Leandro Marziali and Emma Wilson for their help with the initial mouse colony maintenance and sample collection and for providing us with the preliminary data that this study was build on.

## SUPPLEMENTARY FIGURES

**Figure S1:**
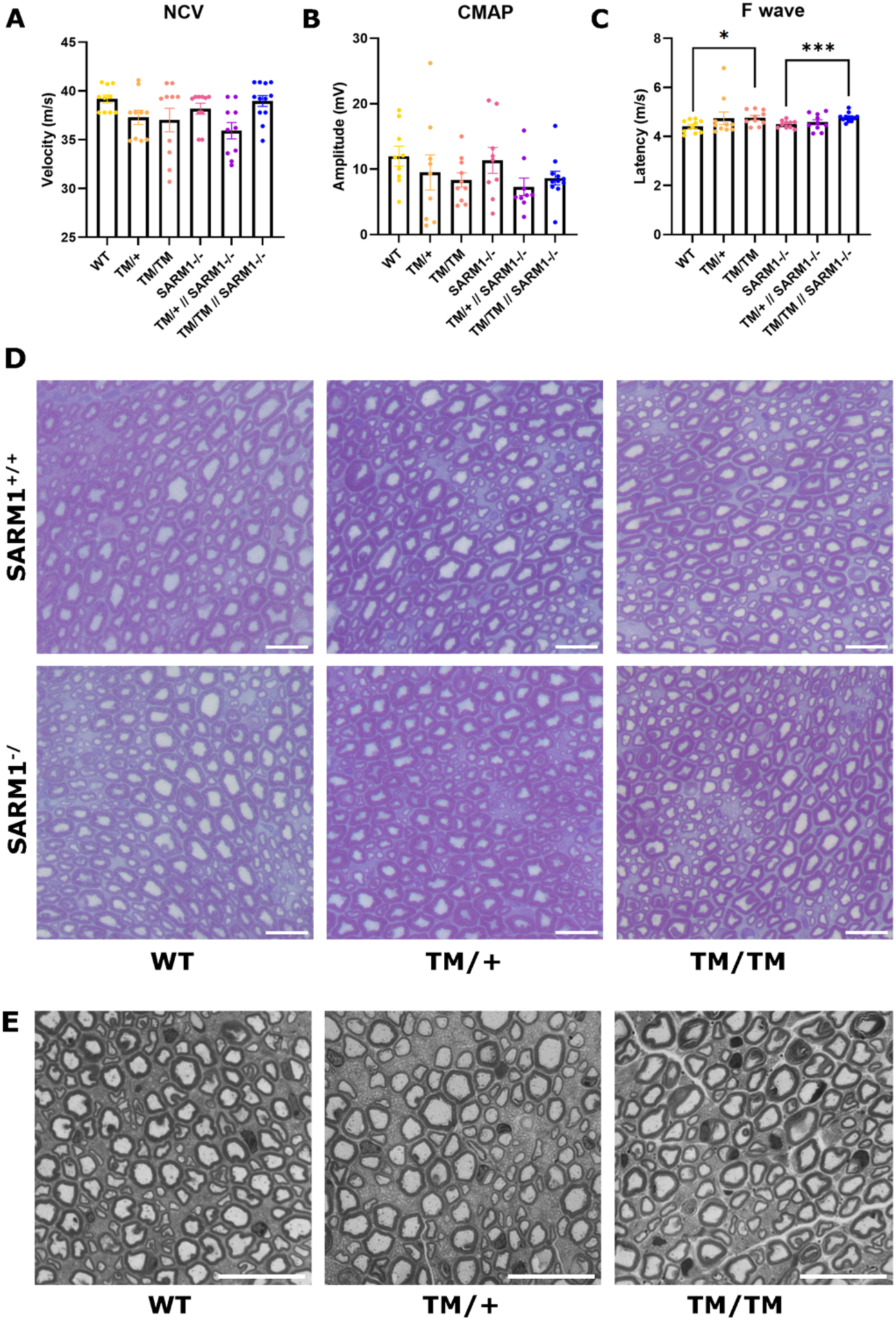
Electrophysiological parameters and peripheral nerve morphology in 4-month-old Mpz^T124M^Sarm1^-/-^ mice. (A) Nerve conduction velocity (NCV), (B) amplitude of compound muscle action potentials (CMAPs) and (C) latency of the F wave were measured at 4 months of age. n = 5 mice per genotype. ***p<0.001; *p<0.05 by one-way ANOVA with Tuckey’s post hoc. Graphs indicate means ± SEM. (D) Representative images of semithin sections of sciatic nerves and (E) saphenous nerves obtained from 4-month-old mice.

**Figure S2:**
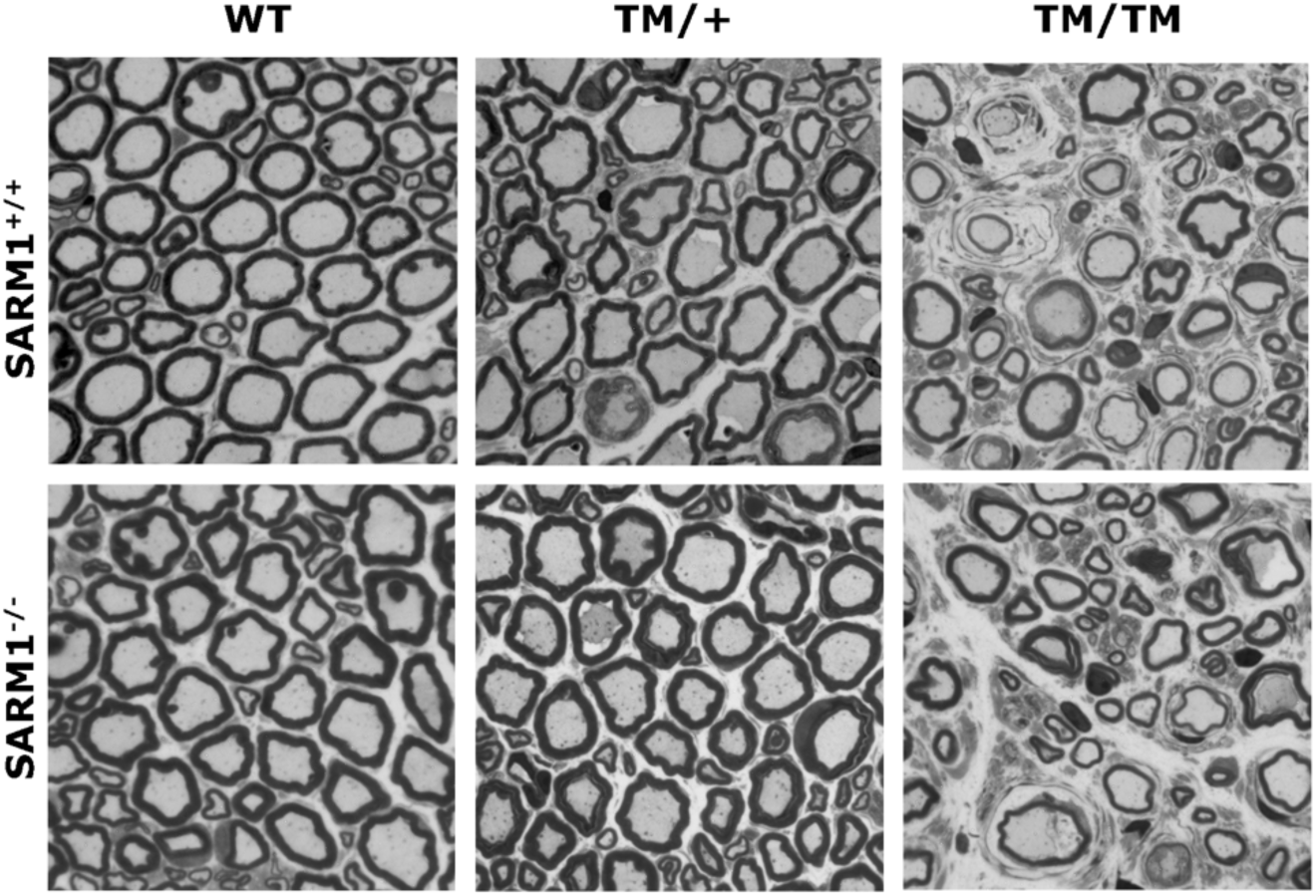
Semithin sections of femoral nerve from 8-month-old Mpz^T124M^Sarm1^-/-^ mice. Representative images of semithin sections of femoral nerves obtained from 8-month-old mice show clear signs of axonal degeneration in homozygous mice.

**Figure S3:**
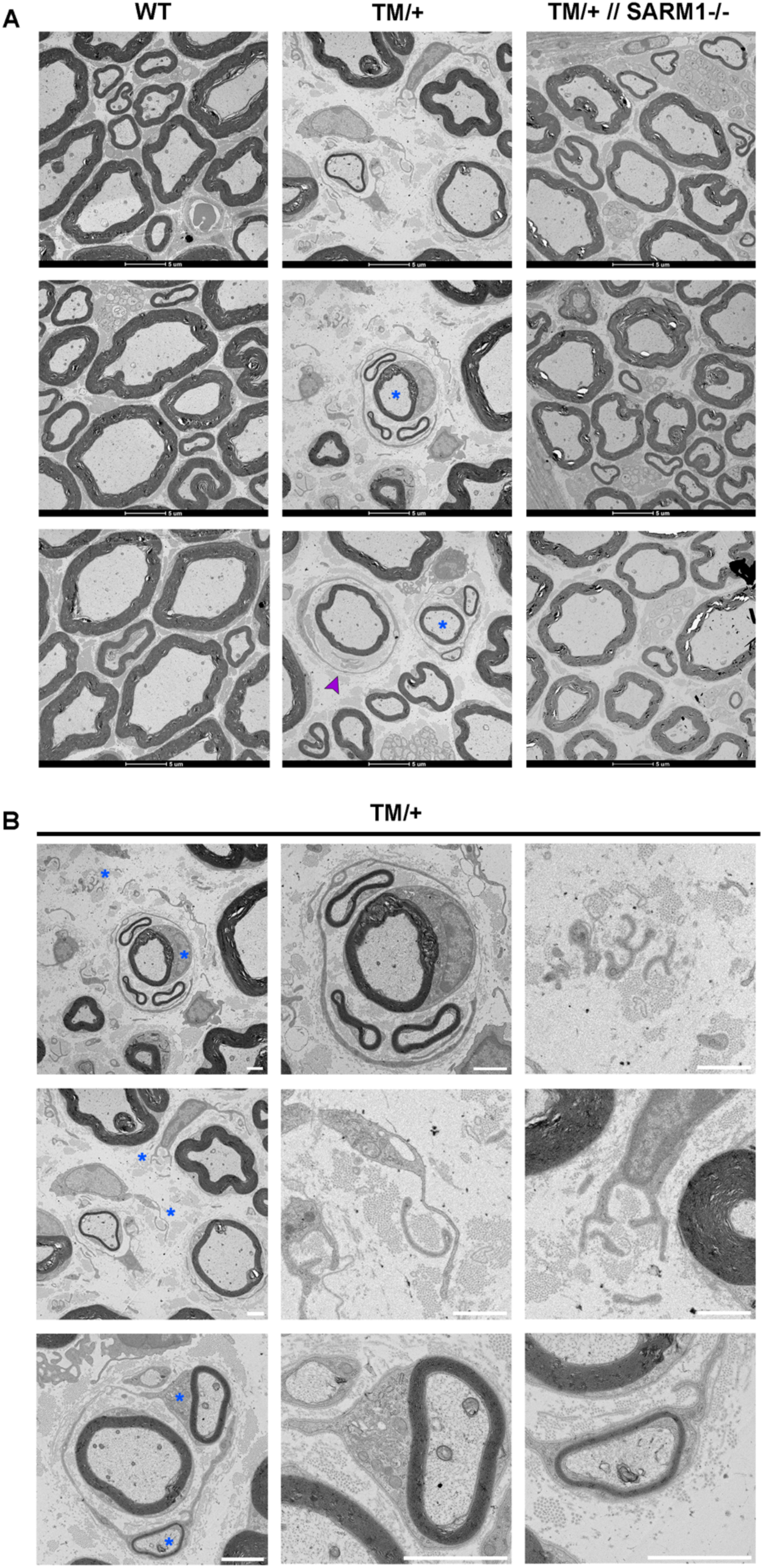
Electron microscopy images of motor femoral nerves from 18-month-old mice. (A) Representative images of electron micrographs from femoral nerves obtained from 18-month-old WT, Mpz^T124M/+^ and Mpz^T124M/+^Sarm1^-/-^ animals. Regenerative clusters are highlighted with a blue asterisks and onion bulb formation with a purple arrowhead. (B) Higher magnification electron micrograph in which each row first shows a larger section of the nerve, and then two magnifications to highlight aberrant figures next to it (magnification sites are indicated by blue asterisks). Scale bars = 2 µm.

**Figure S4:**
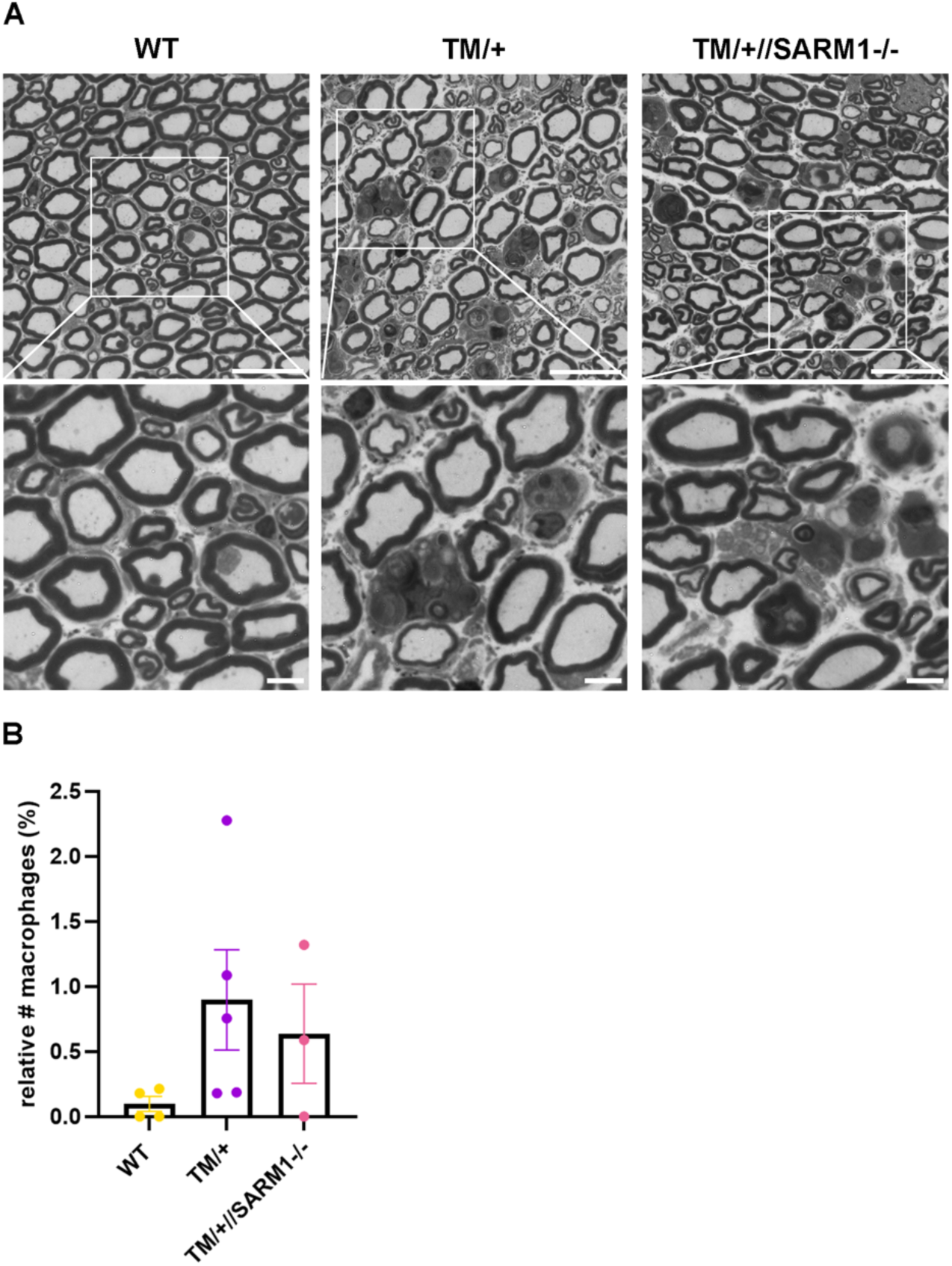
Femoral nerves of 18-month-old mice show phagocytosing macrophages. (A) Representative images of semithin sections of femoral nerves obtained from 18-month-old *WT,* Mpz^T124M/+^ and Mpz^T124M/+^Sarm1^-/-^ animals. The zoom-out indicates phagocytosing macrophages that were quantified in (B). Scale bars = 20µm. Graphs indicate means ± SEM.

## REFERENCES

1. Cashman CR, Höke A. Mechanisms of distal axonal degeneration in peripheral neuropathies. Neurosci Lett. 2015;596:33–50,. 10.1016/j.neulet.2015.01.048

2. Nave K-A, Sereda MW, Ehrenreich H. Mechanisms of Disease: inherited demyelinating neuropathies—from basic to clinical research. Nat Clin Pract Neurol. 2007;3:453–464,. 10.1038/ncpneuro0583

3. Osterloh JM. dSarm/Sarm1 is required for activation of an injury-induced axon death pathway. Science. 2012;337:481–484,. 10.1126/science.1223899

4. Stefano M. A rise in NAD precursor nicotinamide mononucleotide (NMN) after injury promotes axon degeneration. Cell Death Differ. 2015;22:731–742,. 10.1038/cdd.2014.164

5. Loreto A. Mitochondrial impairment activates the Wallerian pathway through depletion of NMNAT2 leading to SARM1-dependent axon degeneration. Neurobiol Dis. 2020;134:104678,. 10.1016/j.nbd.2019.104678

6. Figley MD. SARM1 is a metabolic sensor activated by an increased NMN/NAD^+^ ratio to trigger axon degeneration. Neuron. 2021;109:1118–1136 1111,. 10.1016/j.neuron.2021.02.009

7. Gerdts J, Brace EJ, Sasaki Y, DiAntonio A, Milbrandt J. SARM1 activation triggers axon degeneration locally via NAD^+^ destruction. Science. 2015;348:453–457,. 10.1126/science.1258366

8. Essuman K. The SARM1 Toll/Interleukin-1 Receptor Domain Possesses Intrinsic NAD^+^ Cleavage Activity that Promotes Pathological Axonal Degeneration. Neuron. 2017;93:1334–1343 1335,. 10.1016/j.neuron.2017.02.022

9. Henninger N. Attenuated traumatic axonal injury and improved functional outcome after traumatic brain injury in mice lacking Sarm1. Brain. 2016;139:1094–1105,. 10.1093/brain/aww001

10. Geisler S. Prevention of vincristine-induced peripheral neuropathy by genetic deletion of SARM1 in mice. Brain. 2016;139:3092–3108,. 10.1093/brain/aww251

11. Sato-Yamada Y. A SARM1-mitochondrial feedback loop drives neuropathogenesis in a Charcot-Marie-Tooth disease type 2A rat model. J Clin Invest. 2022;132. 10.1172/JCI161566

12. Bloom AJ, Mao X, Strickland A, Sasaki Y, Milbrandt J, DiAntonio A. Constitutively active SARM1 variants that induce neuropathy are enriched in ALS patients. Mol Neurodegener. 2022;17:1. 10.1186/s13024-021-00511-x

13. Arthur-Farraj P, Loreto A. Targeting SARM1: from inhibition for neuroprotection to activation for neuroablation. Trends Pharmacol Sci. 2025; 10.1016/j.tips.2025.09.003

14. Shackleford G. A new mouse model of Charcot-Marie-Tooth 2J neuropathy replicates human axonopathy and suggest alteration in axo-glia communication. PLoS Genet. 2022;18:1010477,. 10.1371/journal.pgen.1010477

15. Kim Y. MyD88-5 links mitochondria, microtubules, and JNK3 in neurons and regulates neuronal survival. J Exp Med. 2007;204:2063–2074,. 10.1084/jem.20070868

16. Feng G. Imaging Neuronal Subsets in Transgenic Mice Expressing Multiple Spectral Variants of GFP. Neuron. 2000;28:41–51,. 10.1016/S0896-6273(00)00084-2

17. Touvier T. Activation of XBP1s attenuates disease severity in models of proteotoxic Charcot-Marie-Tooth type 1B. Brain. 2025; 10.1093/brain/awae407

18. Sandelius Å. Plasma neurofilament light chain concentration in the inherited peripheral neuropathies. Neurology. 2018;90:518–524,. 10.1212/wnl.0000000000004932

19. Sasaki Y, Engber TM, Hughes RO, Figley MD, Wu T, Bosanac T, et al. cADPR is a gene dosage-sensitive biomarker of SARM1 activity in healthy, compromised, and degenerating axons. Exp Neurol. 2020;329:113252. 10.1016/j.expneurol.2020.113252

20. Fazal SV. SARM1 detection in myelinating glia: sarm1/Sarm1 is dispensable for PNS and CNS myelination in zebrafish and mice. Front Cell Neurosci. 2023;17:1158388,. 10.3389/fncel.2023.1158388

21. Bertini A, Gentile L, Cavallaro T, Tozza S, Saveri P, Russo M, et al. Phenotypic spectrum of myelin protein zero-related neuropathies: a large cohort study from five mutation clusters across Italy. J Neurol Neurosurg Psychiatry. 2024;96:47–53. 10.1136/jnnp-2024-333842

22. Kemshead JT, Ritter MA, Cotmore SF, Greaves MF. Human Thy-1: expression on the cell surface of neuronal and glial cells. Brain Res. 1982;236:451–461,. 10.1016/0006-8993(82)90727-2

23. Verdú E, Butí M, Navarro X. Functional changes of the peripheral nervous system with aging in the mouse. Neurobiol Aging. 1996;17:73–77,. 10.1016/0197-4580(95)02010-1

24. Walsh ME. Use of Nerve Conduction Velocity to Assess Peripheral Nerve Health in Aging Mice. J Gerontol Ser A. 2014;70:1312–1319,. 10.1093/gerona/glu208

25. Klein D. Macrophage Targeting Protects Nerve Structure and Improves Muscle Innervation in a Mouse Model of Charcot-Marie-Tooth 2J. Glia. 2025;73:2369–2385,. 10.1002/glia.70074

26. Arthur-Farraj PJ, Latouche M, Wilton DK, Quintes S, Chabrol E, Banerjee A, et al. c-Jun Reprograms Schwann Cells of Injured Nerves to Generate a Repair Cell Essential for Regeneration. Neuron. 2012;75:633–47. 10.1016/j.neuron.2012.06.021

27. Jonghe P. The Thr124Met mutation in the peripheral myelin protein zero (MPZ) gene is associated with a clinically distinct Charcot-Marie-Tooth phenotype. Brain. 1999;122:281–290,. 10.1093/brain/122.2.281

28. Hanemann CO, Gabreëls-Festen AA, Jonghe P. Axon damage in CMT due to mutation in myelin protein P0. Neuromuscul Disord. 2001;11:753–756,. 10.1016/s0960-8966(01)00229-2

29. Evgrafov OV, Mersiyanova I, Irobi J, Van Den Bosch L, Dierick I, Leung CL, et al. Mutant small heat-shock protein 27 causes axonal Charcot-Marie-Tooth disease and distal hereditary motor neuropathy. Nat Genet. Nature Publishing Group; 2004;36:602–6. 10.1038/ng1354

30. Pan X, Xie J, Li Z, Xiang Y, Yu Y, Cai Q, et al. Stress-Driven Selective Neuronal Vulnerability in Charcot–Marie–Tooth Disease: From Prodromal Pathology to Therapeutic Implications. Cells. 2026;15:271. 10.3390/cells15030271

31. Bosanac T. Pharmacological SARM1 inhibition protects axon structure and function in paclitaxel-induced peripheral neuropathy. Brain. 2021;144:3226–3238,. 10.1093/brain/awab184

32. Kurihara S. Axonal and demyelinating forms of the MPZ Thr124Met mutation. Acta Neurol Scand. 2003;108:157–160,. 10.1034/j.1600-0404.2003.00110.x

33. Dingwall CB, Strickland A, Yum SW, Yim AKY, Zhu J, Wang PL, et al. Macrophage depletion blocks congenital SARM1-dependent neuropathy. J Clin Invest [Internet]. American Society for Clinical Investigation; 2022 [cited 2026 June 24];132. 10.1172/JCI159800

34. Collins JM. Sarm1 knockout modifies biomarkers of neurodegeneration and spinal cord circuitry but not disease progression in the mSOD1G93A mouse model of ALS. Neurobiol Dis. 2022;172, 105821. 10.1016/j.nbd.2022.105821

35. Moss KR. SARM1 knockout does not rescue neuromuscular phenotypes in a Charcot-Marie-Tooth disease Type 1A mouse model. J Peripher Nerv Syst. 2022;27:58–66,. 10.1111/jns.12483

36. Hatton CL. Genetic Deletion of Sarm1 in Mouse Models of Three Neurological Diseases. J Peripher Nerv Syst. 2025;30:70052,. 10.1111/jns.70052

37. Rice AD, Tadenev ALD, Hines TJ, Funke JR, Burgess RW. SARM1 Inhibition in Three Mouse Models of Charcot-Marie-Tooth Disease. J Peripher Nerv Syst. 2025;30:70053,. 10.1111/jns.70053

